# Neuronal microstructural changes in the human brain are associated with neurocognitive aging

**DOI:** 10.1101/2024.01.11.575206

**Authors:** Kavita Singh, Stephanie Barsoum, Kurt G Schilling, Yang An, Luigi Ferrucci, Dan Benjamini

**Affiliations:** Multiscale Imaging and Integrative Biophysics Unit, National Institute on Aging, NIH, Baltimore, MD, USA; Department of Radiology and Radiological Sciences, Vanderbilt University Medical Center, Nashville, TN, USA; Brain Aging and Behavior Section, National Institute on Aging, NIH, Baltimore, MD, USA; Translational Gerontology Branch, National Institute on Aging, NIH, Baltimore, MD, USA

**Keywords:** microstructure, gray matter, aging, MRI, diffusion, neurocognitive aging

## Abstract

Gray matter (GM) alterations play a role in aging-related disorders like Alzheimer’s disease and related dementias, yet MRI studies mainly focus on macroscopic changes. Although reliable indicators of atrophy, morphological metrics like cortical thickness lack the sensitivity to detect early changes preceding visible atrophy. Our study aimed at exploring the potential of diffusion MRI in unveiling sensitive markers of cortical and subcortical age-related microstructural changes and assessing their associations with cognitive and behavioral deficits. We leveraged the Human Connectome Project-Aging cohort that included 707 unimpaired participants (394 female; median age = 58, range = 36–90 years) and applied the powerful mean apparent diffusion propagator model to measure microstructural parameters, along with comprehensive behavioral and cognitive test scores. Both macro- and microstructural GM characteristics were strongly associated with age, with widespread significant microstructural correlations reflective of cellular morphological changes, reduced cellular density, increased extracellular volume, and increased membrane permeability. Importantly, when correlating MRI and cognitive test scores, our findings revealed no link between macrostructural volumetric changes and neurobehavioral performance. However, we found that cellular and extracellular alterations in cortical and subcortical GM regions were associated with neurobehavioral performance. Based on these findings, it is hypothesized that increased microstructural heterogeneity and decreased neurite orientation dispersion precede macrostructural changes, and that they play an important role in subsequent cognitive decline. These alterations are suggested to be early markers of neurocognitive performance that may distinctly aid in identifying the mechanisms underlying phenotypic aging and subsequent age-related functional decline.

## Introduction

Magnetic resonance imaging (MRI) macrostructural volumetric studies focused on the gray matter (GM) have demonstrated consistent correlations between older age and longitudinal decline of cortical volume and thickness (Fjell et al., 2009; Resnick et al., 2003; Sowell et al., 2003). Though, spatial localization and degree of atrophy are not homogeneous across the aging brain (Fjell et al., 2014), and for example, frontal and temporal lobes exhibit the highest degree of atrophy, parietal lobe exhibits moderate changes, whereas the occipital lobe appears to remain relatively intact (Peters, 2006). At the macroscopic level, the occurrence and extent of age-related volumetric changes of white matter (WM) show variability and inconsistency (Gunning-Dixon et al., 2009). However, diffusion MRI (dMRI), which probes meso- and microstructural information (Callaghan, 2011), enables investigation of age-related changes in individual fiber pathways of the brain that are not detected by traditional MRI.

Diffusion tensor imaging (DTI) (Basser et al., 1994), which describes the distribution of diffusion displacements using a simplistic Gaussian model, is the most widely used dMRI experimental and theoretical framework to study microstructural integrity of brain tissue. Studies using DTI have unveiled complex and non-linear age-related patterns in brain tissue during maturation and degeneration, typically reflecting increases in anisotropy and decreases in diffusivity during childhood, adolescence and early adulthood, and subsequent anisotropy decreases, and diffusivity increases in adulthood and senescence (Beck et al., 2021; Madden et al., 2009; Schilling et al., 2022). Nonetheless, contrary to basic assumptions in DTI, diffusion within heterogeneous biological tissues is non-Gaussian, primarily due to the complex microstructure involving the presence of cell membranes, organelles, and distinct liquid compartments (Novikov et al., 2018). Consequently, while DTI exhibits considerable structural sensitivity, it is hindered by these limitations and by its lack of specificity, which in turn impede its interpretation. To enhance specificity, various multicomponent biophysical models have been developed (Assaf & Basser, 2005; Zhang et al., 2012), some of which have been effectively used to study aging (Gozdas et al., 2021; Nazeri et al., 2015; Vogt et al., 2020). While these models may be appropriate to study WM, they present significant limitations in studying GM because they rely on strict assumptions about tissue composition and physical properties (Jespersen et al., 2019; Kundu et al., 2023; Palombo et al., 2020; Veraart et al., 2019). Consequently, to date most dMRI microstructural studies on brain aging have predominantly concentrated on WM. This trend is also evident in studies related to myelin water fraction imaging (Mackay et al., 1994), which have solely centered on age-related alterations in WM (Bouhrara et al., 2020; Faizy et al., 2018).

The importance of developing a sensitive MRI framework to assess age related microstructural changes in GM has been highlighted by recent anti-amyloid-β clinical trials that showed increased GM atrophy was observed despite evidence of target engagement and a slowdown in cognitive decline (Mintun et al., 2021; Swanson et al., 2021). Such outcomes may be attributed to underlying micro- and meso-structural changes, reflecting alterations in axonal connectivity, cellular morphology, and the neuroinflammatory response in GM. Although few studies suggested that dMRI is sensitive to cortical microstructural changes in unimpaired and impaired aging, they were limited to large GM regions and were performed on relatively small cohorts (Gozdas et al., 2021; Nazeri et al., 2015; Vogt et al., 2020). A recent pilot study showed that the mean apparent propagator model (MAP-MRI) model (Özarslan et al., 2013), a dMRI experimental and theoretical framework which does not rely on any biophysical assumptions, can leverage multi-shell dMRI data and provide high sensitivity to age-related microstructural changes in GM (Bouhrara et al., 2023). Echoing these results, cortical microstructural alteration derived from MAP-MRI were shown to be highly sensitive to multiple aspects of the Alzheimer’s disease pathological cascade (Spotorno et al., 2023). Indeed, MAP-MRI may facilitate the study of healthy and diseased white and gray matter as it directly measures the diffusion propagator in each voxel, enabling comprehensive insights into tissue diffusion processes and microstructural parameter derivation, unconstrained by assumptions about tissue compartments that are unlikely to hold in normal aging and disease.

In this work, we use MAP-MRI to probe cerebral GM microstructure and architecture in the large cohort of the Human Connectome Project-Aging (HCP-A). This represents the most in-depth application of MAP-MRI in normal aging to date, incorporating state-of-the-art processing procedures to allow microstructural investigation of cortical and subcortical GM regions. In this study, we aim to: (1) Characterize GM macro- and microstructural differences with age using volumetric and MAP-MRI, respectively; (2) Investigate associations of GM macro- and microstructure with cognitive measures and determine whether one or both can be used as predictors of neurobehavioral performance.

## Results

Imaging data from 707 subjects aged 36 to 90 years were included in this study, along with their available behavioral and cognitive test scores (Bookheimer et al., 2019). The list of tests and respective scores stratified by participants age is provided in Table 1. A total of 56 cortical and subcortical GM regions of interest (ROIs), covering most of the brain, were included in the study (Fig. 1). For each MRI metric and ROI, only statistically significant associations that survived multiple comparison correction were shown (*p*_FDR_<0.05), where *β*_age_ expresses the rate of change from young to adult (at sample’s mean age of 59.6 years), *β*_age_^2^ reflects the direction and steepness of the quadratic association, and *β*_MR_ reflects the rate of change of the various MR metrics with respect to the different neurobehavioral test scores. Further details about the statistical models can be found in the Methods section.

**Figure 1:**
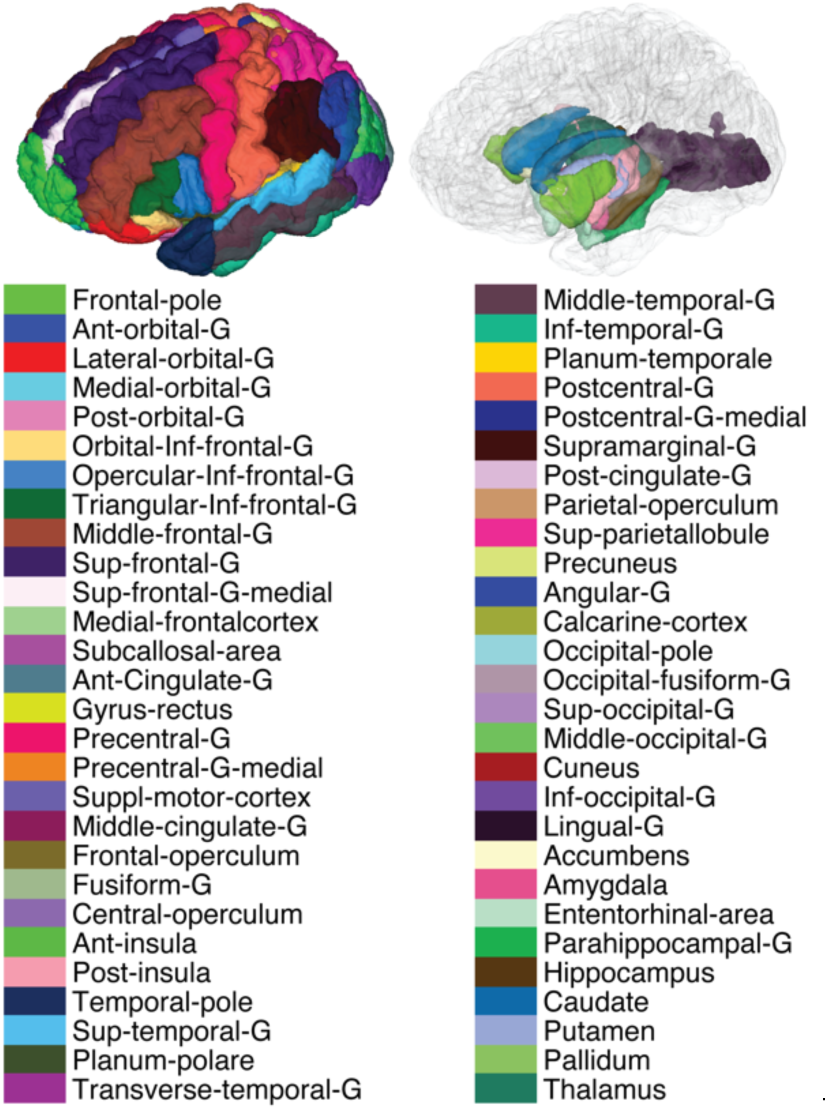
Cortical and subcortical view of the 56 brain regions of interest (ROI) used in the study. Note that the ROI-based analysis was performed in the native space of each subject to minimize errors resulting from interpolation. Abbreviations: Ant-anterior; G-gyrus; Post-posterior; Inf-inferior; Sup-superior.

**Table 1.** Participant demographics (N = 707) and list of neurobehavioral test scores expressed as mean (standard deviation). YOE: years of education; PSM: Picture Sequence Memory; RAVLT: Rey Auditory Verbal Learning Test.

|  | All | Age ≤ 60 | Age > 60 | *p*-value | missing samples | Neurobehavioral domain |
| --- | --- | --- | --- | --- | --- | --- |
| **N** | 707 | 383 | 324 |  |  | - |
| **Age [years]** | 59.7 (14.9) | 47.9 (7.0) | 73.6 (8.3) | <0.001 | 0 | - |
| **Sex, F/M** | 394/313 | 221/162 | 173/151 | 0.251 | 0 | - |
| **YOE [years]** | 17.5 (2.2) | 17.4 (2.2) | 17.6 (2.1) | 0.116 | 1 | - |
| **Flanker** | 8.00 (0.82) | 8.26 (0.73) | 7.69 (0.82) | <0.001 | 86 | Executive function, attention |
| **PSM** | 505 (103) | 551 (92.7) | 448 (86.0) | <0.001 | 85 | Episodic memory |
| **Trail Making A** | 29.9 (11.8) | 26.3 (10.1) | 34.1 (12.3) | <0.001 | 5 | Cognitive processing speed |
| **Trail Making B** | 74.1 (41.9) | 64.5 (34.6) | 85.4 (46.9) | <0.001 | 3 | Executive functioning |
| **RAVLT 1** | -0.32 (0.47) | -0.26 (0.41) | -0.40 (0.52) | <0.001 | 10 | Episodic memory (retroactive) |
| **RAVLT 2** | -0.15 (0.60) | -0.08 (0.50) | -0.24 (0.69) | <0.001 | 7 | Episodic memory (proactive) |

\**p*-values were derived from two sample t-test for continuous variables and chi-square test for categorical variables.

The MAP-MRI framework does not rely on any assumptions and is suitable to use in complex microstructures, offering consequential advantages over biophysical model-based dMRI methods. While MAP-MRI provides five distinct microstructural metrics, namely, return to the axis/origin/plane probabilities – RTAP, RTOP, RTPP (jointly referred to as the zero-displacement probabilities), the propagator anisotropy (PA), and the non-Gaussianity (NG), they do not yield straightforward structural metrics and requires interpretation. These variables can be thought of as proxies of cellular density and extracellular volume (zero-displacement probabilities), microstructural shape and neurite orientation dispersion (PA), and microstructural heterogeneity (NG).

### Linear associations between age and regional GM macro- and microstructure

We first examined the linear associations with age (at centered age of 59.6 years) of several MRI parameters expressed by the *β*_age_ in our regression model. Macro- and microstructure in GM were captured by ROI volume, and MAP-MRI parameters, respectively (Fig. 2). After correction for multiple comparisons, 55, 51, 42, 41, 45, and 48 out of the 56 cortical and subcortical ROIs for NG, PA, RTAP, RTOP, RTPP, and Volume, respectively, were significantly associated with age. Microstructurally, most of the regions showed positive associations of NG and PA and negative associations of RTAP, RTOP, and RTPP and age. However, some regions, for example the medial frontal cortex, gyrus rectus, accumbens, caudate and putamen, exhibited opposing trends of the RTAP, RTOP, and RTPP metrics. Macrostructurally, negative correlation of ROI volume and age was generally observed.

**Figure 2:**
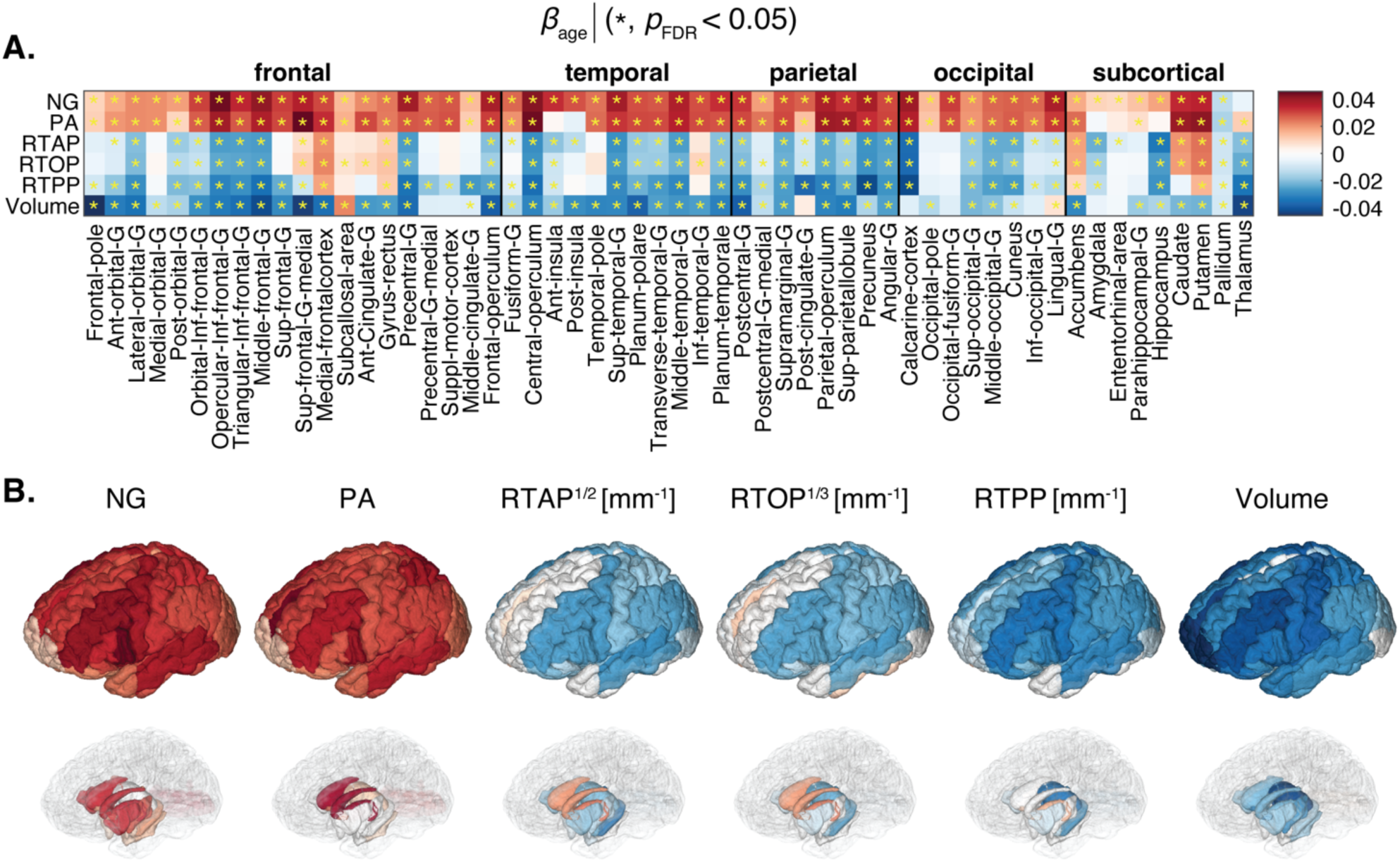
Linear associations of MAP-MRI and volumetric metrics and age. (A) The *β*_age_ coefficients are shown as a matrix for MAP-MRI and volumetric z-normalized features across all 56 ROIs. Blocks marked with an asterisk (*) represent associations meeting the *p*_FDR_<0.05 threshold. (B) 3D visualization of significant results in cortical (top) and subcortical (bottom) view.

### Quadratic associations between age and regional GM macro- and microstructure

Nonlinear age relationships in GM are characterized by the direction and steepness of the quadratic curvature (*β*_age_^2^), as depicted in Fig. 3. However, not all GM regions survived multiple comparisons correction, and specifically, NG and PA did not display significant quadratic age relationships in most ROIs. The prominent negative associations observed with RTAP, RTOP, and RTPP, which are sensitive to cellular density and extracellular volume, were primarily concentrated in frontal, parietal, and temporal regions, including the lateral orbital gyrus, posterior-orbital gyrus, anterior insula, medial temporal gyrus, planum temporale, supramarginal gyrus, and parietal operculum, as well as in the medial temporal gyrus, entorhinal area, parahippocampus, hippocampus, and amygdala. While occipital regions yielded fewer significant results, the angular gyrus and calcarine cortex stood out. Importantly, subcortical regions and memory-related areas such as the hippocampus, known for their vulnerability in age-related cognitive decline disorders, demonstrated significant inverted U-shape associations with age. Macrostructurally, inverted U-shape quadratic correlation of ROI volume and age was observed in the frontal, parietal, temporal, subcortex, and limbic system regions.

**Figure 3:**
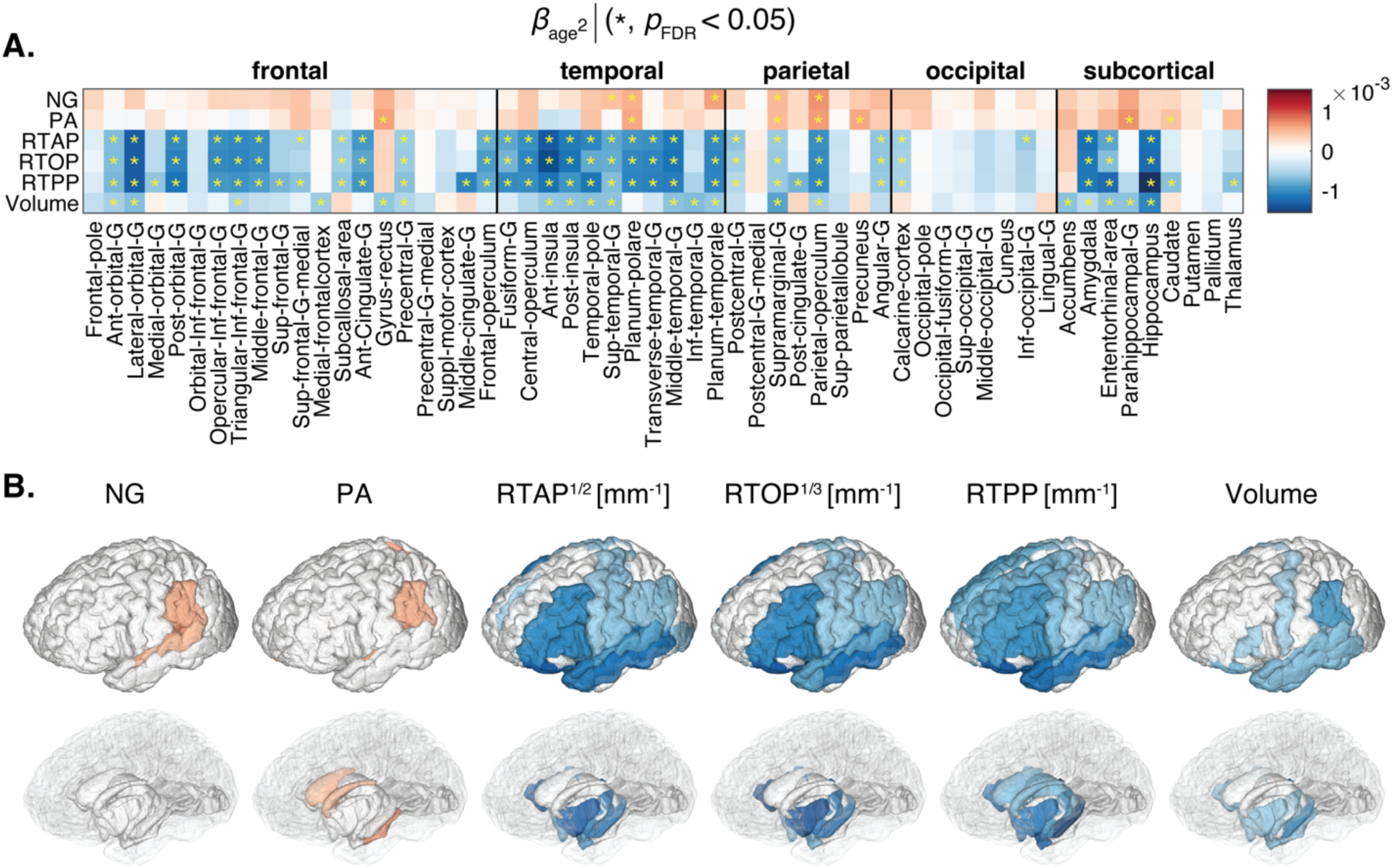
Quadratic associations of MAP-MRI and volumetric metrics and age. (A) The *β*_age_^2^ coefficients are shown as a matrix for MAP-MRI and volumetric z-normalized features across all 56 ROIs. Blocks marked with an asterisk (*) represent associations meeting the *p*_FDR_<0.05 threshold. (B) 3D visualization of significant results in cortical (top) and subcortical (bottom) view.

### Macro- and microstructural age trajectories

The peak ages for each MR metric within each ROI are illustrated in Fig. 4, omitting peak ages over 80 years and negative values. As anticipated, the lack of significant quadratic age associations for NG and PA in most ROIs is consistent with the absence of plausible peak age values in some of those regions. When considering peak age as an indicator of onset of age-related changes, we observed diverse trends.

**Figure 4:**
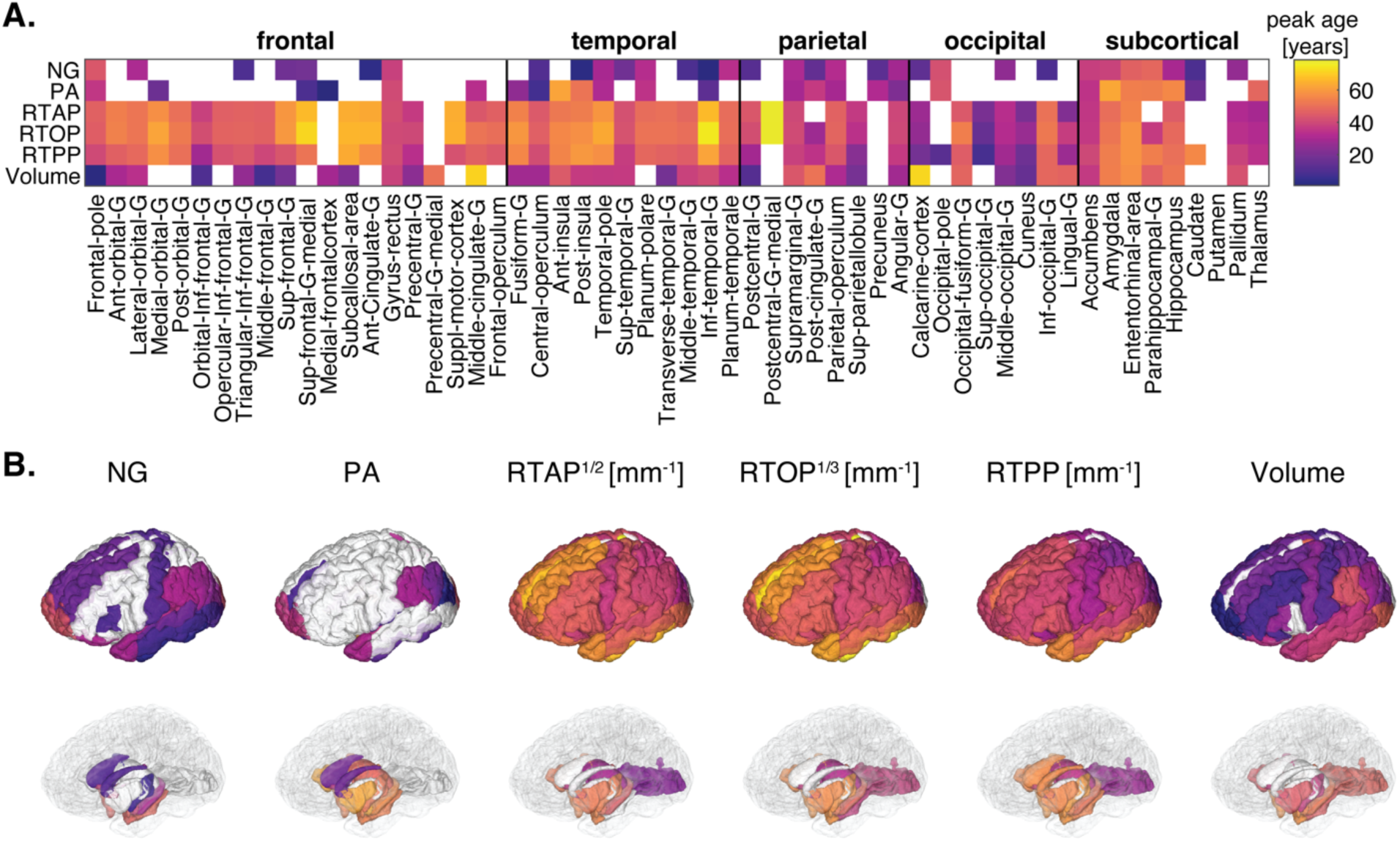
Peak ages for MAP-MRI and volumetric within each ROI. (A) Matrix representation of peak age across all 56 ROIs. (B) 3D visualization of significant peak ages in cortical (top) and subcortical (bottom) view.

Microstructurally, MAP-MRI metrics revealed varying peak ages across different brain regions, indicating the spatial distribution of age-related changes, as well as their onset. The NG and PA generally showed the earliest peaks, which could point to early onset of increased microstructural heterogeneity and reduced neurite orientation dispersion, respectively. On the other hand, the zero-displacement probabilities exhibited considerably later peak ages in the late forties, indicating that reduced cellular density and increased extracellular volume may exacerbate at a later stage in life. Macrostructurally, peak age with respect to volumetric changes in most cortical regions demonstrated later onset compared with NG and PA, and earlier onset compared with the zero-displacement probabilities. Memory-related subcortical regions like the parahippocampus and hippocampus peaked at a relatively older age (about 50 years).

Further evidence supporting NG and PA as early markers of age-related microstructural changes are given by examining their linear associations with age specifically within a younger subset of the cohort (age<50 years, N=227). This analysis showed significant associations, exclusively with NG and PA, in numerous brain regions (Supplementary Fig. 1).

### Micro- and not macrostructure is associated with neurobehavioral performance

We proceeded to examining the relationship between the macro- and microstructural MRI metrics and cognitive test scores by modeling them as outcomes, adjusted for age, sex and education. Figure 5 shows the linear association of different MR metrics as expressed by *β*_MR_ with respect to the Flanker, PSM, Trail Making, and RAVLT tests scores. Interestingly, almost exclusively the NG and PA metrics resulted in significant correlations with the various cognitive and behavioral test scores that survived multiple comparisons correction. These metrics, which relate to microstructural heterogeneity and to neurite orientational dispersion, respectively, were also found to exhibit the earliest onset of age-related changes. The NG appeared to be the most robust MAP-MRI predictor towards cognitive performance, and therefore its correlation coefficient with respect to the tests, *β*_NG_, is shown in sagittal, coronal, and axial views in Fig. 6 to provide visualization of its spatial distributions.

**Figure 5:**
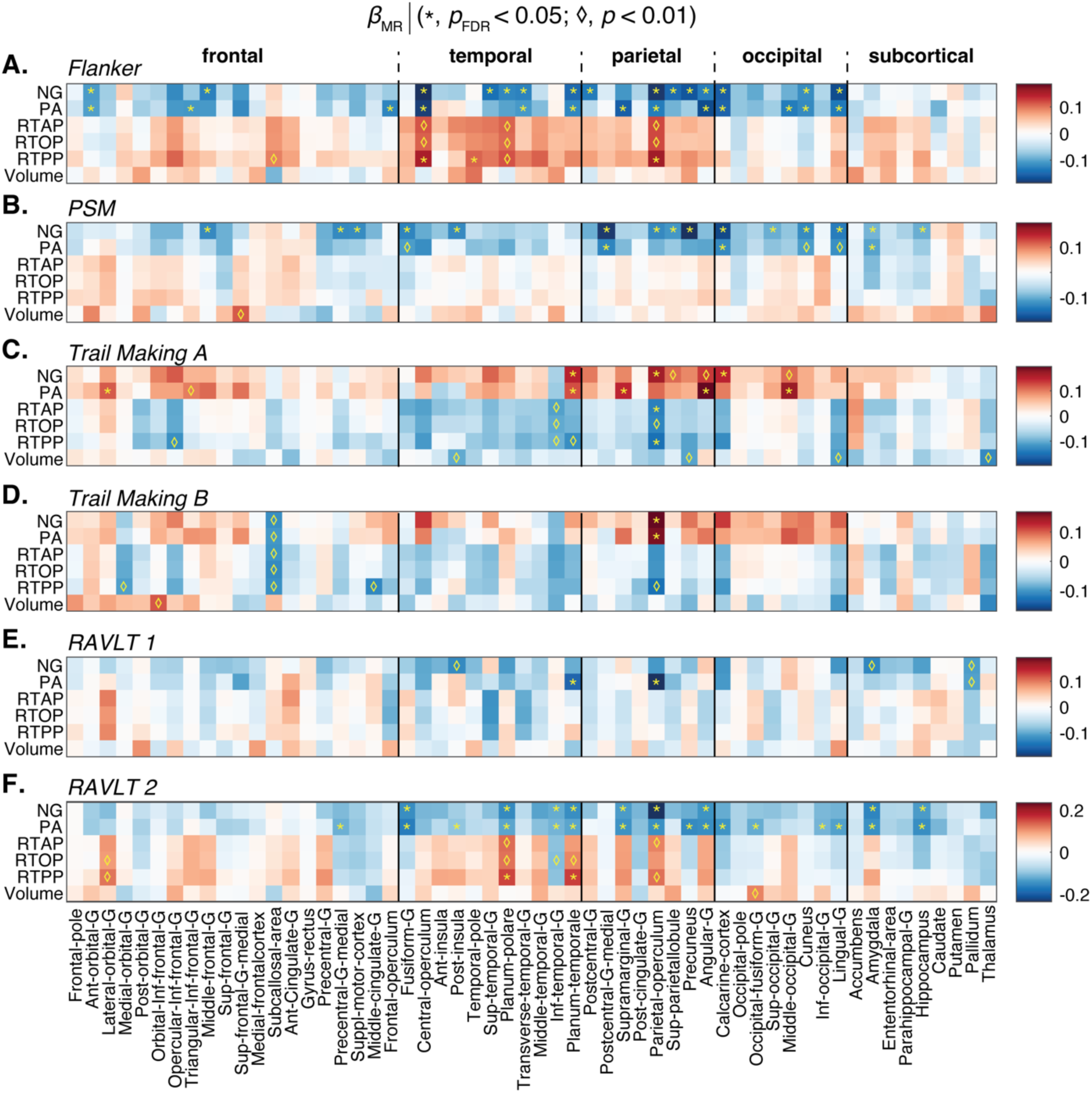
Significant results of second statistical model examining the relationship between the macro- and microstructural MRI metrics and five cognitive test scores by modeling them as outcomes, adjusted for age, sex and years of education. Significant associations are expressed using the regression parameters *β*_MR_. Blocks marked with an asterisk (*) represent associations meeting the *p*_FDR_<0.05 threshold. Blocks marked with a diamond (♢) represent associations meeting the *p*<0.01 threshold without FDR correction. For all tests, the lower the score the worse the performance, except for the Trail Making task, in which the opposite is true. PSM; Picture Sequence Memory test; RAVLT: Rey auditory verbal learning test.

**Figure 6:**
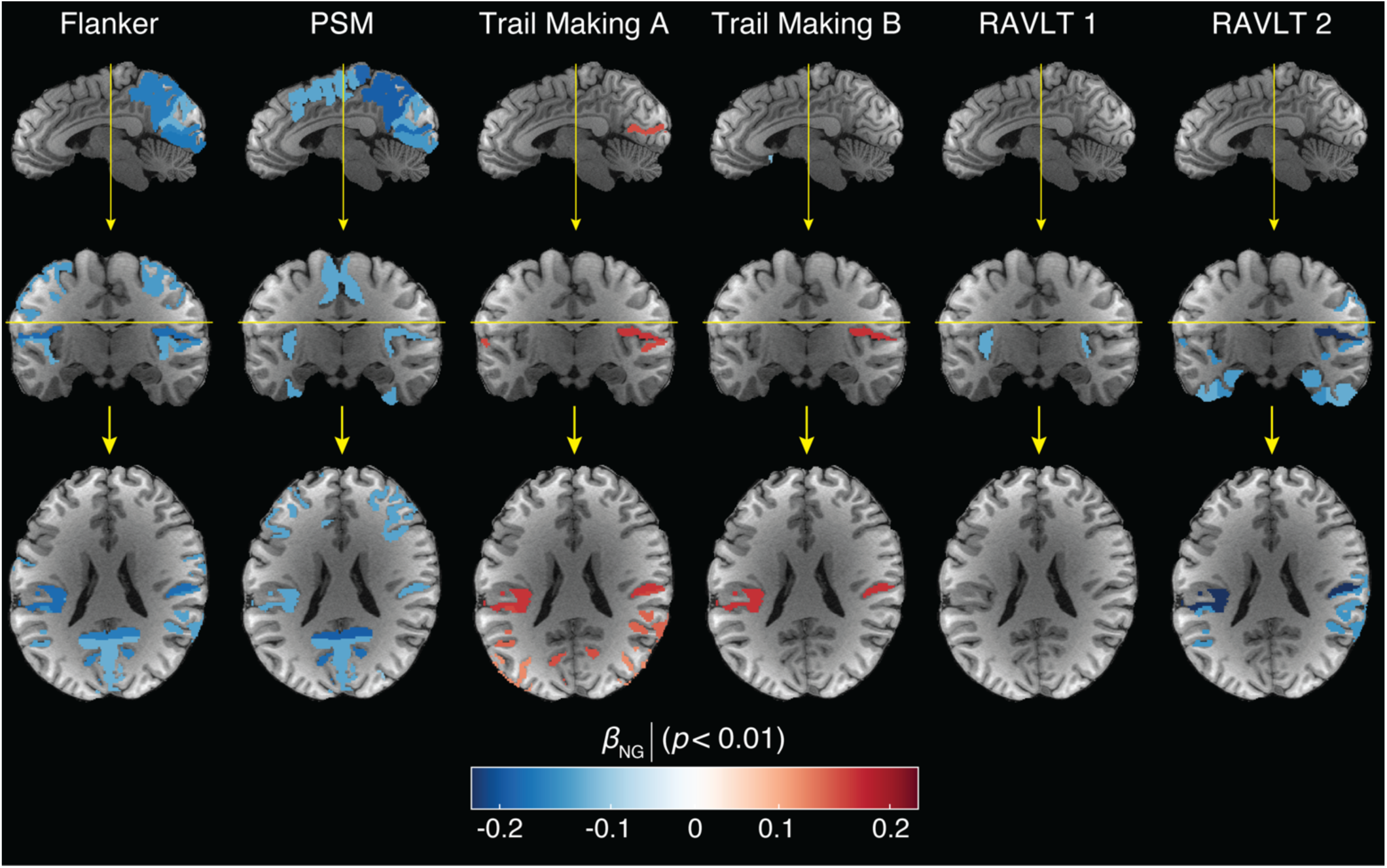
Axial, coronal, and sagittal views of significant results of relationship between the non-Gaussianity (NG) and 6 cognitive test scores. Age-related significant associations are expressed using the regression parameters *β*_NG_ with *p* < 0.01. For all tests, the lower the score the worse the performance, except for the Trail Making task, in which the opposite is true. PSM: Picture Sequence Memory test; RAVLT: Rey auditory verbal learning test. Yellow lines indicate sectional planes.

The Flanker inhibitory control task, a measure of executive function and attention where lower scores indicate poorer performance, is known to exhibit functional associations with the cingulo-opercular and fronto-parietal control networks (Dosenbach et al., 2007). Our findings reveal significant negative correlations of both NG and PA metrics in the anterior orbital gyrus, central and parietal operculum, transverse temporal gyrus, planum temporale, calcarine cortex, lingual gyrus, and cuneus, as visualized in Figs. 5A and 6A. Positive associations with the zero displacement probabilities were observed in the central and parietal operculum.

The PSM task score assesses episodic memory, in which the lower the score the worse the performance, exhibited significant negative correlations with NG and PA in brain regions responsible for encoding new learning into episodic memory, which encompassed primary sensory areas (visual, auditory, somatosensory) and memory-related regions (prefrontal cortex, amygdala, hippocampus), as shown in Figs. 5B and 6. Notably, significant negative correlations were observed in several brain regions, including frontal, temporal, parietal and occipital cortical regions, as well as the amygdala and hippocampus.

The Trail Making tests evaluate task completion time, examining psychomotor speed, visuospatial search, and target-directed motor tracking that are linked to prefrontal and parietal cortices (Varjacic et al., 2018), as well as links to functional connectivity between right frontoparietal and visual networks (Küchenhoff et al., 2021). As such, the higher the score the worse the performance. In our study, the Trail Making A test significantly correlated with NG and PA in network regions like lateral orbital gyrus, planum temporale, angular gyrus (related to attention, self-processing), and visual network regions including middle occipital gyrus, depicted in Figs. 5C and 6. Trail Making B test scores, which measure executive functioning, were significantly correlated with NG and PA in the parietal operculum.

The RAVLT test, with variants 1 and 2, assesses episodic memory function. Lower scores indicate poorer performance. RAVLT 1 relates to retroactive interference and was negatively correlated with PA in planum temporale and parietal operculum, areas linked to auditory processing (Tanaka & Kirino, 2018) and multimodal integration (Mălîia et al., 2018), respectively. RAVLT 2 relates to proactive interference and showed widespread correlations with PA, NG, and RTPP in proactive interference mediating regions involved in memory (Hamilton et al., 2022), including the posterior parietal cortex, medial-temporal lobe, occipital regions, amygdala, and hippocampus, involved in semantic memory and working memory.

Importantly, macrostructural volumetric information was not predictive of neurobehavioral performance in any of the tests.

## Discussion

Gray matter alterations have been shown to be associated with aging and various aging-related disorders such as Alzheimer’s, Parkinson’s, and dementias. Examinations of aging-related brain changes using MRI have predominantly focused on the macroscopic level in GM or employed DTI to study WM microstructure. With the advent of advanced models like MAP-MRI, we can now study microstructural changes in GM as well. By conducting an extensive analysis of a large, cross-sectional dataset from the Human Connectome Project encompassing 707 subjects aged between 36 and 90 years, we examined MAP-derived micro- and volumetric-derived macrostructural features, in addition to cognitive and behavioral metrics, across 56 GM brain regions, and their relationships with age. Notably, while volumetric MRI did not display associations to age-related cognitive and behavioral performance, MAP-MRI features exhibited strong correlations with memory and executive functions in the corresponding GM regions. Building upon our prior study, in which the superiority of MAP-MRI over DTI in assessing GM was demonstrated (Bouhrara et al., 2023), we presented here an in-depth and comprehensive characterization of age-related microstructural alterations in GM using MAP-MRI across the adult lifespan and elucidated their associations with cognitive and behavioral performance.

The most significant advantage of the MAP-MRI framework over biophysical model-based dMRI methods is that it operates without assumptions and is suitable to use in complex microenvironments. While MAP-MRI metrics do not provide straightforward structural metrics, we can suggest their biophysical interpretation. The zero-displacement probabilities (RTAP, RTOP, RTPP) are inversely relate to the nominal size of microstructures and are affected by cellular density and extracellular volume. The PA captures the three-dimensional directionality of diffusion, and in GM it encompasses information about microstructural shape and neurite orientation dispersion. Finally, the NG quantifies the deviation from DTI’s simple tensor model and reflects the heterogeneity of microstructures.

### Gray matter macro- and microstructural changes with age: linear and non-linear associations

When examining the age-related linear and quadratic trends of MAP-MRI features, we identified positive relationships with PA and NG, and negative correlations with zero-displacement probabilities in most GM regions (see Figs. 2 and 3). Our findings established that the associations between NG and PA with age in GM were primarily linear, while the zero-displacement probabilities displayed substantial quadratic associations in most GM regions, except for the occipital regions. Additionally, we observed macrostructural changes involving volumetric reductions in most regions. Much like the results from MAP-MRI, the volumetric reductions in occipital regions did not exhibit significant quadratic associations with age.

The negative quadratic associations with age trends we observed with respect to the zero displacement probabilities are in agreement with a previous MAP-MRI pilot study (Bouhrara et al., 2023), and also with the widely observed behavior of diffusivity metrics in GM across the adult lifespan (Salminen et al., 2016). These results can be hypothesized to indicate reduced cellular density and increased extracellular volume in GM with age. In addition, local inflammation, which is a common feature of aging brain, leads to activated state of microglia and astrocytes in the aged brain (Mattson & Arumugam, 2018). The resulting glial morphological change, in particular astrocytes, which includes hypertrophy of cell body and stem processes (Sofroniew, 2020), is therefore hypothesized to contribute as well to the decrease of MAP-MRI measured zero displacement probabilities with age.

The PA provides a higher-order assessment of DTI’s fractional anisotropy (FA) and quantifies the directional dependence of the diffusion process. From our analysis, we found that the PA is positively linearly associated with age in almost all GM regions. While age-related patterns of increased anisotropy during childhood, adolescence and early adulthood, and subsequent anisotropy decrease in adulthood and senescence (Resnick et al., 2003; Schilling et al., 2022), have been observed in WM, increased FA with age in deep brain GM structures has been previously reported (Pfefferbaum et al., 2010). Further, older age was associated with reduced neurite orientation dispersion in widespread cortical regions (Bouhrara et al., 2023; Gozdas et al., 2021). Thus, microstructurally, our findings with respect to PA are hypothesized to reflect morphological changes in neurons, which are expected to decrease in number, shorten, and become less branched with fewer spines (Dickstein et al., 2013).

The zero-displacement probabilities and the PA extend the DTI model and its derived parameters, providing metrics that could be more sensitive to subtle pathological events and more suitable to complex microstructure (Avram et al., 2016a; Saleem et al., 2021). On the other hand, the NG provides completely distinct information that quantifies the dissimilarity between the full propagator and its Gaussian component, and in essence reflects the deviation from DTI’s tensor model. With that, while the NG is the least interpretable MAP-MRI parameter, it carries unique information and also seems to be the most microstructurally sensitive with respect to the aging brain (Bouhrara et al., 2023). Similar to the PA in our study, we found significant positive linear associations of the NG with age in almost all GM regions. The non-Gaussianity of the diffusion processes can be driven primarily by restriction (Benjamini & Basser, 2019), membrane permeability (Williamson et al., 2019), and cellular degradation (Benjamini et al., 2020), which together form microstructural heterogeneity. The robust increases of NG in normative aging that we observed supports the microstructural scenario of dendritic changes (Dickstein et al., 2007), reactive astrocytes, especially hypertrophy of stem processes, and increased membrane permeability due to myelin alterations through processes like demyelination and remyelination (J. Lee & Kim, 2022).

Some zero-displacement probabilities associations in subcortical regions exhibited an opposing trend. Specifically, RTAP, RTOP, and RTPP in the accumbens, caudate, and putamen demonstrated positive linear correlations with age, in agreement with previous MAP-MRI findings (Bouhrara et al., 2023). Despite declines in memory and cognitive functions, numerous studies have noted heightened prefrontal cortex activity in healthy older adults (Davis et al., 2008). This phenomenon has been ascribed to either a compensatory functional shift from posterior to anterior regions, aiding cognitive performance despite posterior cortical impairment, or a primary age-related functional decline in the prefrontal cortex, resulting in non-specific increased activity (Nyberg et al., 2012; Park et al., 2004). In our study, the increases in zero-displacement probabilities with age coupled with positive NG and PA age associations in these regions may be indicative of reduced extracellular volume and relative resilience to glial reactivity. However, these hypotheses should be specifically examined in future studies.

### Trajectories of macro- and microstructural changes with age are heterogeneous

Although impossible to determine without a longitudinal study, information about the period at which age-related micro- and macrostructural changes become apparent can be gleaned from estimating the peak age with respect to each MR parameter (Fig. 4). The peak age can therefore be used as an indicator of onset of age-related changes and help elucidate the trajectories of different micro- and macrostructural changes. Our results showed diverse trends, in which some changes appear to precede others. Specifically, we found that the NG and PA reach peak age earliest, followed by macroscopic volume, and finally zero-displacement probabilities. These results imply that increased microstructural heterogeneity and decreased neurite orientation dispersion (reflected from NG and PA, respectively) may precede macrostructural changes (reflected from regional volumes) and changes in cellular density and extracellular volume (reflected from RTAP, RTOP, and RTPP). Additional support for NG and PA as early indicators of age-related microstructural changes is provided by analyzing their linear correlations with age specifically within a younger subset of the cohort, which revealed significant associations (Supplementary Fig. 1).

Interestingly, regions associated with the limbic system, paralimbic areas, and limbic-related regions like the anterior cingulate, amygdala, entorhinal area, parahippocampus, hippocampus, temporal pole, and cortex-insula, exhibited the latest peak age based on both MAP-MRI and volumetric analyses. These findings align with the concept of the limbic network displaying relative age resilience, given its role in emotions and memories. For instance, the limbic network is recognized for demonstrating the least susceptibility to age-related changes in cortical thickness in young healthy adults (Bajaj et al., 2017), minimal volumetric reductions in cognitively healthy aging populations (Fujita et al., 2023), and relatively stable DTI parameters in limbic-related white matter tracts, such as the parahippocampal cingulum (Bennett et al., 2015) or rostral and dorsal cingulum (Stadlbauer et al., 2008). These studies consistently report the preservation, relative stability, and sparing of the medial temporal lobe in healthy aging, encompassing the entorhinal and parahippocampal cortices across all age groups. Our findings here align with these studies, demonstrating preserved micro- and macrostructural characteristics in the limbic network. It is hypothesized that cognitive reserve mechanisms might safeguard against changes in the micro- and macrostructure of the brain within the limbic network, a topic that warrants exploration in future research.

### Gray matter microstructure linked to neurobehavioral performance

Significant associations between performance in various behavioral and cognitive tasks and microstructural changes, primarily reflected from NG and PA, were observed across several brain regions (Figs. 5 and 6). Importantly, while we showed that GM volumetric data is highly correlated with age, these macroscopic changes proved to be poor predictors of neurobehavioral function. These findings, combined with the earlier peak age of the NG and PA compared with volumetric changes, can be hypothesized to reflect that microstructural precede macrostructural changes, and that the former can predict the development of the latter. These results underscore the importance of quantifying microstructural and architectural features in GM in the context of aging.

Specifically for the Flanker task, significant negative correlations of NG and PA were identified in regions crucial for executive function (fronto-parietal control network) and attention (cingulo-opercular network). The former exhibits late development and early decline, aligning with established literature (Craik & Bialystok, 2006). Inhibition, one of the key processes of executive functions that is controlled by inferior frontal gyrus (Aron et al., 2003), and was seen as significant negative association with PA in our results as well. Hasher and Zacks (Hasher et al., 1999) proposed that inhibition is specifically impaired with advanced adult age and age-related inhibition deficit has been reported in other studies (Andrés et al., 2008). The cingulo-opercular network, which supports cognitive control and visual processing speed, has been reported to exhibit reduced structural and functional connectivity in aging (Ruiz-Rizzo et al., 2019). These reports correspond well with the observed reductions in neurite orientation dispersion reflected from the negative correlations of the Flanker score with PA.

Our study also revealed wide-spread associations of PSM task scores and NG and PA parameters. Episodic memory function is reported to be extremely sensitive to cerebral aging (Nyberg et al., 2012). Episodic memory, initially linked with the hippocampus, is now known to be a multi-component network including the medial temporal lobe (entorhinal, perirhinal, and parahippocampal cortices), prefrontal cortex, lateral temporal neocortex, and posterior parietal regions (Dickerson & Eichenbaum, 2010). We observed negative associations between NG and PA values, not just in regions linked to memory (such as the prefrontal cortex, amygdala, hippocampus, etc.) but also in primary sensory areas (including visual, auditory, and somatosensory regions). These findings are in line with the established ‘network’ and ‘events’ involved in encoding episodic memory, where first the elements that constitute an event register across the primary sensory areas. These inputs from the primary cortices are then projected to unimodal association areas, and subsequently to polymodal (prefrontal, posterior, and limbic) association areas. Lastly, these inputs are integrated and maintained for short term (Petrides, 1995), or consolidated for long-term (medial temporal structures) (McGaugh, 2000).

The Trail Making A and B tests provide information on visual search, scanning, speed of processing, mental flexibility, and executive functions, and have been shown to decline with age based on neuropsychological studies (Tombaugh, 2004). However, the brain regions involved during this task specific activity and its differential performance in healthy older adults has only been recently confirmed using functional MRI studies (Talwar et al., 2020a). Similar studies using functional near-infrared spectroscopy have also reported parietal, temporal, and occipital regions to be mostly implicated in decline during this task in older adults (Hagen et al., 2014). Specifically, precuneus, angular gyri, hippocampus, inferior parietal lobe and temporal lobe have been reported (Talwar et al., 2020b). These regions of default mode network are suppressed during cognitive stimulation, including performance of complex tasks (e.g., trail making task) and might be the reason of our current findings as well.

The RAVLT tool is widely used for the cognitive assessment of memory consolidation in normal aging, pre-dementia and dementia conditions (Moradi et al., 2017). To allow interpretation, we computed the RAVLT 1 and 2 scores, which are indicative of retroactive and proactive interferences, respectively. The RAVLT 1 associations with PA were found only in the planum temporale and parietal operculum, which are involved in auditory processing and as an integration center within a multimodal network (Mălîia et al., 2018), respectively. Although disentangling these two phenomena is beyond the scope of current work, we did find memory regions significantly associated with the RAVLT 2 score, e.g., amygdala and hippocampus, which could be explored further in future work. Various facets of encoding, consolidation, and retrieval processes are hypothesized to involve the precuneus, suggesting the utilization of visual imagery during encoding (Gonsalves et al., 2004), involvement of the posterior medial temporal lobe and visual areas linked to the formation of true memories (Kim & Cabeza, 2007), and the auditory cortex, as evidenced in both the literature (Straube, 2012) and our present study.

Despite the study’s strengths, it has limitations. The large number of subjects is beneficial, but they were mainly white, highly educated individuals from the HCP-A database, limiting generalization due to potential biases related to race, ethnicity, culture, and education. Being cross-sectional, the study lacks a longitudinal design, but future analysis of the forthcoming HCP-A follow-up data could enhance its scope. A longitudinal design would allow to directly determine the order in which micro- and macrostructural changes takes place across the lifespan, and whether earlier changes can predict the development of later changes and cognitive decline. The age cohort, starting from 36 years, restricts exploration of changes across the age spectrum, potentially emphasizing late maturation and degeneration phases. Additionally, while the HCP-A dMRI dataset provides good orientational coverage and AP-PA encoding, the maximal b-value is 3000 s/mm^2^. The relatively low diffusion sensitization in our study limits our ability to capture subtle features of the underlying diffusion propagators. Consequently, we measured the propagators using 22 coefficients associated with the most significant basis functions in a MAP-MRI series expansion truncated at order 4 (Özarslan et al., 2013). The inclusion of higher b-values, up to 6000 s/mm^2^ (Huang et al., 2021), would have allowed us to consider terms up to order 6, as found to be optimal in clinical applications (Avram et al., 2016b). Subsequent studies ought to explore and delineate microscopic alterations in the shape, dimensions, and geometry of cerebral tissue throughout the lifespan. While the conventional linear diffusion encoding scheme that is used here does not allow the decoupling of size and microscopic orientation, utilizing multiple diffusion encoding schemes (Benjamini et al., 2012; Mitra, 1995) such as planar or spherical (Topgaard, 2017) is recommended to study these effects. Additionally, the simultaneous application of relaxation and diffusion encoding should be considered in future investigations (Benjamini et al., 2021, 2023; Martin et al., 2021).

In conclusion, this comprehensive analysis utilizes a large cross-sectional dataset and applies the advanced MAP-MRI model to delineate GM microstructural relationships with age and neurobehavioral performance across healthy aging. Notably, while we did not identify macrostructural volumetric associations with neurobehavioral scores, we did, for the first time, establish robust correlations between GM microstructure and age-related neurobehavioral performance. Based on our findings, we hypothesize that increased microstructural heterogeneity and decreased neurite orientation dispersion precede macrostructural changes, and that they play an important role in subsequent cognitive decline. This study provides valuable insights that could assist in early differentiation of cognitively healthy aging from pathological conditions, including mild cognitive impairment, Alzheimer’s, and other dementia-related disorders.

## Methods

### Study design and participants

Data was obtained from Human Connectome Project-Aging (HCP-A) Lifespan 2.0 Release. It included 725 healthy adults aged 36 to 90+ years old acquired across four acquisition sites using matched MRI scanning protocols (Harms et al., 2018). All participants were screened for causes of cognitive decline and exhibited typical health for their age without stroke, clinical dementia. The study was approved and monitored by the Institutional Review Board. Written informed consent was obtained from all participants in the study. After written consent, the Montreal Cognitive Assessment (MoCA) was administered, and participants meeting the determined normal threshold for their age bracket were considered eligible for the study. No participants were excluded based on medication use, although self-reported medication use was recorded during the study visit to investigate or avoid specific medication confounds. Essential health assessments known to show associations to brain circuitry during typical aging, as well as in dementia and other diseases, was performed and has been previously described (Bookheimer et al., 2019). This process excluded participants who have been diagnosed and treated for major psychiatric disorders (e.g., schizophrenia, bipolar disorder) or neurological disorders (e.g., stroke, brain tumors, Parkinson’s Disease).

Of 725 healthy subjects’ data available, we included 707 subjects for MAP-MRI and volumetric analysis based on quality control of the raw data. The study population demographics is provided in Table 1. During HCP-A 2.0 release, data from participants over the age of 90 years were lumped together due to data policy, and therefore were excluded from our study. Years of education were also recorded and used in the final analysis.

### Imaging data acquisition

To meet the HCP-A recruitment and diversity goals, data were acquired at four different institutions. In all four sites, data were acquired using a 3T scanner (MAGNETOM Prisma, Siemens Healthcare AG, Erlangen, Germany) with a 32 channel head coil.

Structural MRI images were acquired using T1- and T2-weighted (T1W and T2W) contrasts. T1-weighted imaging was performed using a multi-echo MPRAGE sequence with 0.8 mm isotropic voxel size, TR = 2500 ms, TI = 1000 ms, TE = 1.8/3.6/5.4/7.2 ms, and flip angle of 8°. T2-weighted imaging was performed using a T2w-SPACE protocol with 0.8 mm isotropic voxel size, TR = 3200 ms and TE = 56.4 ms. Both sequences used embedded volumetric navigators for prospective motion correction and selective reacquisition of the lines in k-space corrupted by motion. The mean image of just the first two echoes from the MPRAGE acquisition was used as the input to subsequent processing.

Diffusion-weighted images (DWIs) were acquired using a pulsed gradient spin-echo sequence with 1.5 mm isotropic voxel size and TR/TE = 3230/89.5 ms. Diffusion encoding was acquired with two shells of 1500 and 3000 s/mm^2^ (98-99 directions per shell), and with 28 b-value = 0 s/mm^2^ images interleaved. All datasets were acquired with two phase encoding directions: anterior to posterior (AP), and reversed phase encoding direction (PA). More detailed image acquisition protocols of HCP-A can be found in Harms *et al* (Harms et al., 2018).

### Behavior and cognitive testing

The HCP-A conducted detailed behavioral and cognitive assessments using NIH toolbox (Gershon et al., 2013). The list of tests and respective scores stratified by participants demographics is provided in Table 1. Briefly, the tests included in current study are: 1) Flanker Inhibitory Control and Attention Test, which is an assessment of inhibitory control and attention. The participant is asked to focus on a particular stimulus while inhibiting attention to the stimuli flanking it. 2) Picture Sequence Memory (PSM) test, which assess episodic memory. Participants are shown several pictures, and then asked to reproduce the sequence of pictures as it was presented to them. 3) The Trail Making Test is a neuropsychological test of visual attention and task switching, in which Trail Making A (connecting numbers sequentially) measures cognitive processing speed, whereas Trail Making B (connecting numbers and letters sequentially) measures executive functioning. 4) For a more comprehensive assessment of episodic memory, a widely used neuropsychological measure, the Rey Auditory Verbal Learning Test (RAVLT) was used. The RAVLT consists of a semantically unrelated word list (List A) and a similar interference list (List B). Participants are given five trials to immediately recall as many words as possible from list A of fifteen unrelated words. On the next trial, subjects are asked to immediately recall words of interference list (List B). For the final trial (Trial 6 for list A), the participant is asked to recall as many words as they can from List A. To interpret these scores, we computed RAVLT 1, which is the relative difference between trial 6-list A and trial 5-list A, and RAVLT2, which is the relative difference between list B and trial 1-list A. Low RAVLT 1 score indicative of either high degrees of forgetting during the short delay or retroactive interference, and low RAVLT 2 score indicative of a high degree of proactive interference.

### Data processing

#### Preprocessing

Each participant DWIs were manually quality checked before and during each processing step. The preprocessing modules used in this work are part of the TORTOISE dMRI processing package (Irfanoglu et al., 2023). Briefly, the dMRI data initially underwent denoising with the MPPCA technique (Veraart et al., 2016), which was followed by Gibbs ringing correction (Kellner et al., 2016) for partial k-space acquisitions (H.-H. Lee et al., 2021). Motion and eddy currents distortions were subsequently corrected with TORTOISE’s DIFFPREP module (Rohde et al., 2004) with a physically-based parsimonious quadratic transformation model and a normalized mutual information metric. For susceptibility distortion correction, a T2W image was fed into the DRBUDDI (Irfanoglu et al., 2015) method for phase-encoding distortion correction. The final preprocessed data was output with a single interpolation in the space of an anatomical image at native in-plane voxel size.

#### MAP-MRI parameters estimation

Using these preprocessed DWIs, we estimated the voxel-wise diffusion propagators using a MAP-MRI series expansion truncated at order 4 (Özarslan et al., 2013). MAP-MRI extends the DTI model, harnessing the capabilities of modern dMRI sequences to offer potentially more sensitive metrics for detecting early pathological events in the disease process (Avram et al., 2016a) and to quantify microscopic flow (Benjamini et al., 2019). The commonly derived metrics are the propagator anisotropy (PA), which is a generalized version of DTI’s fractional anisotropy (FA); the non-Gaussianity (NG), which quantifies the dissimilarity between the propagator, and its Gaussian part; and the zero displacement probabilities, including the return to the origin probability (RTOP), the return to the axis probability (RTAP) and the return to the plane probability (RTPP), which comprehensively quantify various features of the three-dimensional diffusion process. We therefore computed images of the MAP parameters: RTOP, RTAP, RTPP, NG, and PA. Note that throughout the paper we report the RTAP^1*/*2^ and RTOP^1*/*3^ values to allow consistency in units of the zero-displacement probability metrics (i.e., 1/mm).

#### Whole brain segmentation and volume estimation

The spatially localized atlas network tiles (SLANT) method was used to perform whole brain segmentation (Huo et al., 2019). Briefly, SLANT employs multiple independent 3D convolutional networks for segmenting the brain. Each of the networks is only responsible for a particular spatial region, thus the task of each network is simplified to focus on patches from a similar portion of the brain. Within this end-to-end pipeline, affine registration, N4 bias field correction, and intensity normalization are employed to roughly normalize each brain to the same space before segmentation. After each network performs its duty, the segmentation labels are fused together to form the labels for 132 anatomical regions based on the BrainCOLOR protocol (https://mindboggle.info/braincolor/). SLANT is publicly available and has shown high intra- and inter-scan protocol reproducibility (Xiong et al., 2019). For this study, we first merged the right and left side brain labels. Out of these labels, we then excluded WM and ventricles regions, and regions that might give non-specific information due to large size (e.g., cerebellum and brainstem). Finally, a total of 56 ROIs were included in the study as shown in Fig. 1. These labels were used, after adjusting for intracranial volume (ICV) to remove the confounding effect of head size, to obtain regional brain volumes. For statistical analysis, SLANT labels were first transformed from T1W space to T2W/DWI space using ANTs rigid registration. Additionally, all ROIs were eroded using a 2×2×2 voxels cubic structuring element to reduce partial volume effects and imperfect image registration and to mitigate structural atrophy seen especially at older ages. The mean NG, PA, RTAP, RTOP, and RTPP values were calculated for each ROI and participant.

#### Statistical analysis

To investigate micro- and macrostructural changes in the brain due to aging, multiple linear regression was applied on the MAP metrics (i.e., NG, PA, RTAP, RTOP, and RTPP) and ROI volume. These six MRI features along with the behavioral and cognitive test scores were z-normalized for further analysis.

We first assessed effect of age on MRI parameters using an age quadratic model, with each mean MRI metric within each ROI as the dependent variable. The model is given by: *P_i_* = *β*0+ *β*_sex_\**sex* + *β*_age_\**age* +*β*_age_^2^\**age*^2^ +*β*_YOE_\**YOE* + *β*_site_\**site* + *β*_inter_\**sex* \**age* + *β*_inter_2\**sex*\**age^2^*, where *P_i_* is the mean ROI value of the parameter of interest (e.g., NG, PA, volume, etc.) of the *i*th ROI. Sex, years of education (YOE), and sex-age interactions were accounted for. In addition, we included a categorical *site* variable as a covariate to account for the four HCP-A data acquisition locations. To estimate the peak age for each MR metric and within each region we equated the first derivative of the quadratic function to zero, such that the peak age was given by -*β*_age_/2*β*_age_^2^.

In our second model, cognitive test scores were used as outcomes, *C* = *β*0+ *β*_sex_*sex + *β*_age_*age +*β*_YOE_\**YOE* + *β*MR* *Pi*, where *C* are the neurobehavioral test scores (i.e., Flanker, PSM, etc.).

We mean centered age variable and used effect coding for sex (0.5 as males and −0.5 as females). These centering methods allow regression coefficient to have meaningful interpretation in the presence of interaction or quadratic terms. False discovery rate (FDR) correction was done to correct for multiple comparisons (Storey, 2002) and the threshold for statistical significance was *p*_FDR_ < 0.05.

Matlab was used for all computations.

## AUTHOR CONTRIBUTIONS

D.B. conceived the study; D.B. and K.S. obtained the datal; K.S., S.B., and K.G.S. analyzed the data; Y.A. designed the statistical models; L.F. contributed to the interpretation of the results and worked on the manuscript; K.S. and D.B. drafted the manuscript. All authors interpreted the findings, commented on the manuscript, and approved the submitted version.

## Acknowledgements

This work was supported by the Intramural Research Program of the National Institute on Aging. KGS was supported by National Institutes of Health award number K01EB032989.

