## Supplementary Fig. 1 for "Neuronal microstructural changes in the human brain are associated with neurocognitive aging"

**Affiliations:**

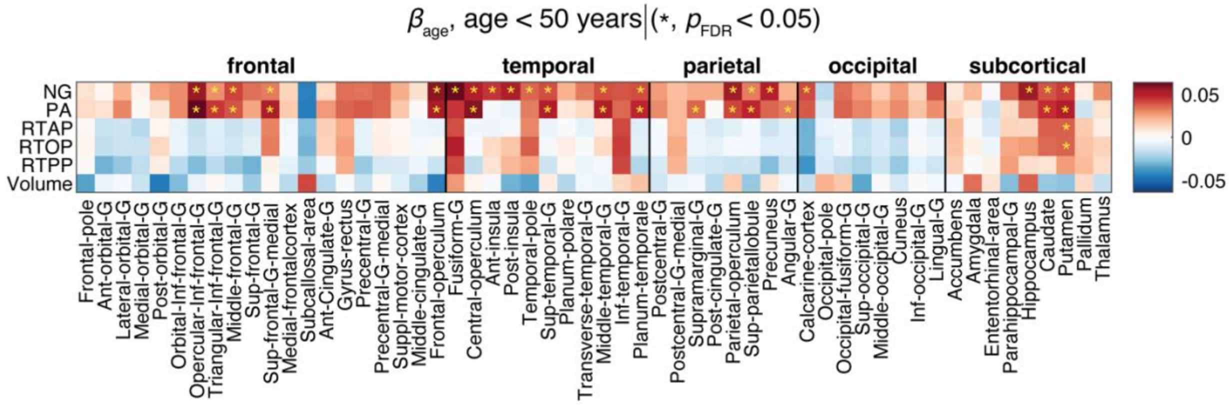


**Supplementary Figure 1**: Linear associations of MAP-MRI and volumetric metrics and age in a subset of subjects under the age of 50 (N=227). The $\beta_{\text{age}}$ coefficients are shown as a matrix for MAP-MRI and volumetric z-normalized features across all 56 ROIs. Blocks marked with an asterisk (*) represent associations meeting the $p_{\text{FDR}}$<0.05 threshold.
